# BK channels of five different subunit combinations underlie the *de novo* KCNMA1 G375R channelopathy

**DOI:** 10.1101/2021.12.22.473917

**Authors:** Yanyan Geng, Ping Li, Alice Butler, Bill Wang, Lawrence Salkoff, Karl L. Magleby

**Affiliations:** Department of Physiology and Biophysics, University of Miami Miller School of Medicine, Miami FL 33136, USA; Department of Neuroscience, Washington University St. Louis, MO 63110, USA; Department of Genetics, Washington University St. Louis, MO 63110, USA; Present address: Department of OBGYN, Washington University St. Louis, MO 63110, USA

## Abstract

The molecular basis of a severe developmental and neurological disorder associated with a *de novo* G375R variant of the tetrameric BK channel is unknown. Here we address this question by recording from single BK channels expressed for a heterozygous G375R mutation. Five different types of functional BK channels were observed: 3% were WT, 12% were homomeric mutant, and 85% were three different types of hybrid channels. All channel types except WT showed a marked gain-of-function in voltage activation and a smaller loss-of-function in single channel conductance, with both becoming more pronounced as the number of mutant subunits per tetrameric channel increased. The molecular phenotype suggested codominance for the two homomeric channels and partial dominance for the hybrid channels. A model in which BK channels are randomly assembled from mutant and WT subunits, with each subunit contributing increments of activation and conductance, approximated the molecular phenotype of the heterozygous G375R mutation.

## Introduction

The BK channel (Slo1, KCa1.1) is a large conductance K^+^ selective channel that is synergistically activated by Ca^2+^ and voltage^1–16^. BK channels are homotetrameric proteins comprised of four large pore-forming (alpha) subunits >1200 amino acids^15^ (Supplementary Fig. 1) encoded by the *KCNMA1* gene. BK channels are widely expressed in many cell types where they modulate smooth muscle contraction^17^, transmitter release^18^, circadian rhythms^19^, repetitive firing^20,21^, and cellular excitability^22^. Mutations in the *KCNMA1* gene that encode the (alpha) subunit of BK channels are associated with a wide range of disease, including epilepsy, dyskinesis, autism, multiple congenital abnormalities, developmental delay, intellectual disability, axial hypotonia, ataxia, cerebral and cerebellar atrophy, bone thickening, and tortuosity of arteries^23–28^.

Studies of the pathogenic properties of BK channels associated with disease have often been incomplete, focusing on the homomeric mutant channels. Yet, for a mutation heterozygous with the wild-type (WT) allele, mutant and WT subunits have the potential to assemble into five different stoichiometries for tetrameric channels with likely differences in functional properties (Fig. 1)^29–33^.

**Fig. 1.**
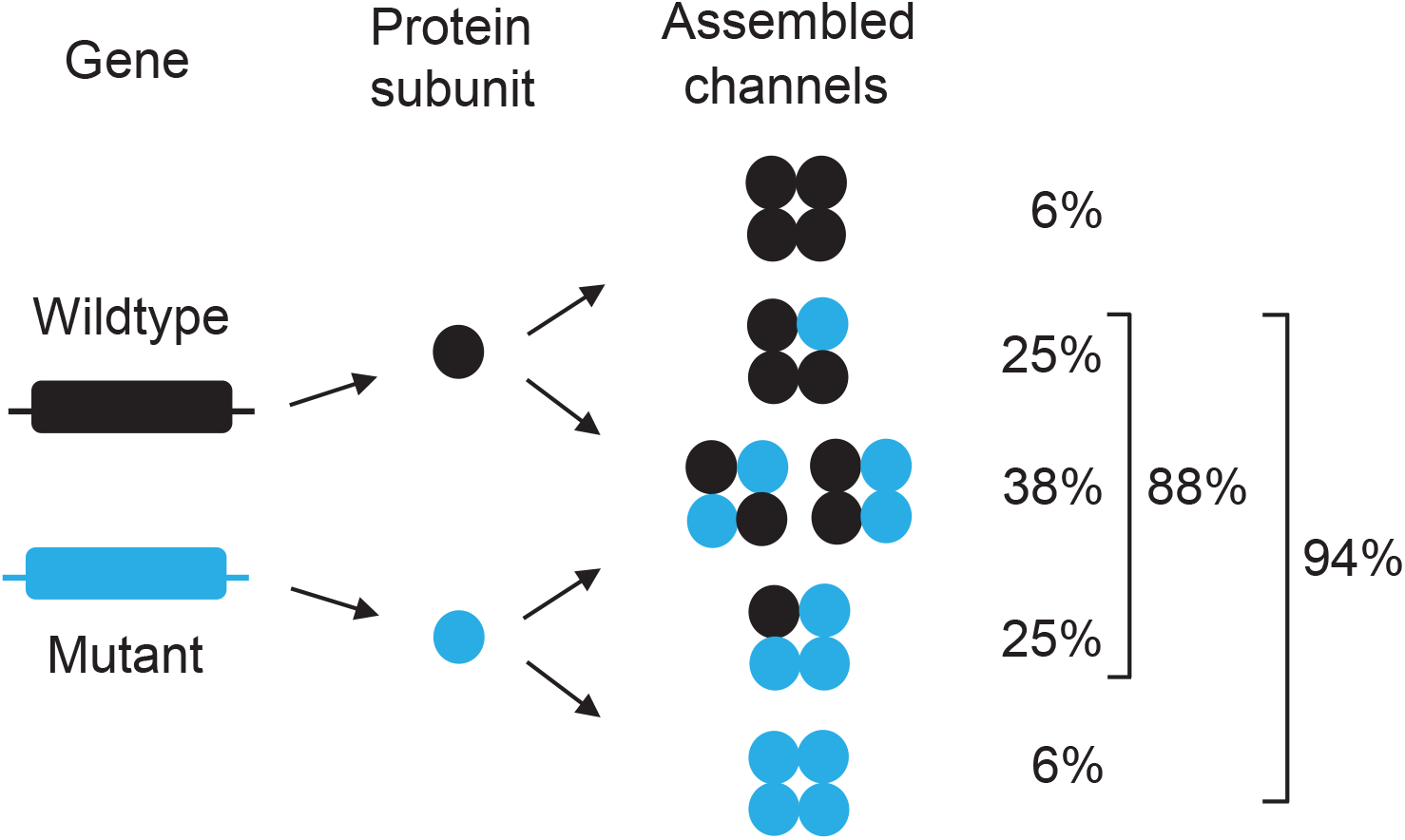
Protein subunits encoded by WT and mutant alleles for a heterozygous mutation could potentially assemble into five different stoichiometries of tetrameric BK channels and six different subunit arrangements^29,30,32,36,55^. In this diagram, WT and mutant alleles (left) encode for WT and mutant protein subunits (middle), which then assemble into tetrameric assembled channels with five different subunit stoichiometries and six different subunit arrangements (right). The listed theoretical frequencies of expression of the five different stoichiometries of assembled channels were calculated with Eqn. 2 in the Methods, which assumes that equal numbers of mutant and WT subunits randomly assemble into tetrameric channels, each with equal probability of reaching the surface membrane. With random assembly, ~6% of the assembled channels are WT comprised of four WT subunits per channel, ~88% are hybrid (mixed subunit) channels with from 1 to 3 mutant subunits per channel, and ~6% are homomeric mutant channels comprised of four mutant subunits per channel. Note that ~94% of the assembled channels can be pathogenic, as they contain from 1 to 4 mutant subunits per channel. The listed percentages of ~6%, ~38%, and ~94% here and throughout the paper have been rounded from the calculated values of 6.25%, 37.5%, and 93.75%. The 37.5% reflects the sum of 25% for hybrid channels with two adjacent mutant subunits and 12.5% for hybrid channels with two diagonal mutant subunits^32^. For experimentation, assembled channels are those channels expressed following injection of a 1:1 mixture of mutant and WT cRNA into *Xenopus* oocytes, to mimic a mutation in which the mutant allele is heterozygous with the WT allele.

Here we show that this is the case for a *de novo* G375R mutation in the (alpha) subunit of BK channels associated with the Liang-Wang Syndrome^25^. Three unrelated children who carried this mutation had a syndromic neurodevelopmental disorder associated with severe developmental delay and polymalformation syndrome^25^. The G375R mutation, located on the back side of the S6 pore-lining helix of the alpha subunit (Supplementary Fig. 1), replaces the hydrogen side chain of glycine with a large arginine side chain that might be expected to distort the subunit structure and gating^34^ as well as add a positive charge that may alter conductance.

To assess the functional effects of a G375R mutation, we recorded currents from whole cells and excised macro patches of membrane after injecting a 1:1 mixture of G375R mutant and WT cRNA encoding for mutant and WT subunits into *Xenopus* oocytes, to mimic a *de novo* mutation heterozygous with respect to the WT allele. When compared to WT, the currents from the 1:1 injection were left shifted more than −120 mV. This large negative shift in activation indicated a pronounced gain-of-function (GOF) mutation at the cellular level, providing a possible explanation for the severity of the disease associated with the heterozygous G375R mutation.

To investigate the molecular basis underlying the cellular response, detailed single-channel recording^35^ after the 1:1 injection suggested that five different types of functional BK channels were expressed: 3% were WT, 12% were homomeric mutant, and 85% were three different types of hybrid (heteromeric) channels. All channel types except WT showed a marked GOF in voltage activation and a smaller LOF in single channel conductance, with both becoming more pronounced as the number of mutant subunits per channel increased. Codominance was observed for the two homotetrameric channels, with homomeric WT channels active at the most positive voltage range of channel activity and homomeric mutant channels active at the most negative. Partial dominance was observed for the three types of hybrid channels, which were activated at voltages intermediate between those of WT and homomeric mutant channels. A model in which BK channels were randomly assembled from mutant and WT subunits, with each subunit contributing increments of activation and conductance, could approximate the molecular phenotype of the heterozygous G375R BK mutation. The possibility that a channelopathy patient with a heterozygous BK channel mutation synthesizes five different types of BK channels in their neurons and other cells, four with aberrant properties, presents a daunting challenge for treatment.

## Results

### G375R subunits left shift BK channel activation

To examine what effect a *de novo* G375R mutation of the pore forming subunit of BK channels would have on channel function when co-expressed with WT subunits, we first compared macroscopic currents flowing through large numbers of WT BK channels expressed following injection of WT cRNA into *Xenopus* oocytes, to those currents expressed following injection of a 1:1 mixture of mutant and WT cRNA to mimic a heterozygous mutation. Injection of a blank without cRNA served as a control. Voltages were stepped from a holding potential of −80 mV to more negative and more positive voltages to reveal the voltage dependent activation of the expressed currents (Fig. 2A; Supplementary Fig. 2). Voltage steps to +50 mV were required to appreciably activate WT BK currents generated from the injection of WT cRNA. In contrast, following a 1:1 injection of G375R mutant and WT cRNA, the expressed BK channels currents were hyperactive, being already active at −140 mV, with depolarization further increasing the response (Fig. 2B, Supplementary Fig. 2). The aberrant BK channels from the 1:1 injection that are responsible for these hyperactive currents could be either homomeric mutant channels, hybrid channels, or both (Fig. 1).

**Fig. 2.**
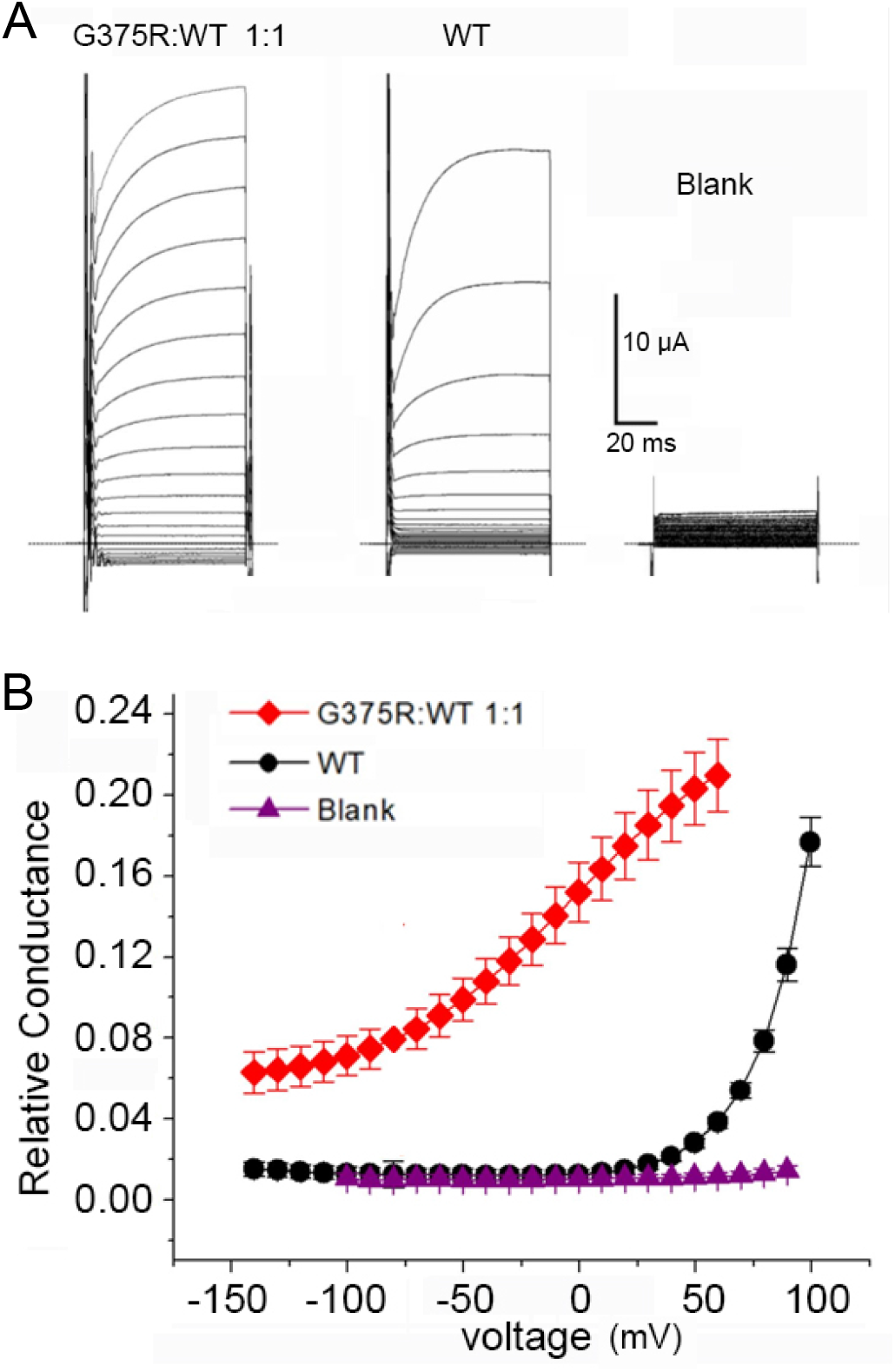
BK channels expressed in oocytes following injection of a 1:1 mixture of G375R mutant and WT cRNA activate at greatly left shifted negative potentials compared to BK channels expressed following injection of only WT cRNA. (***A***) Whole cell currents recorded from oocytes with the two-electrode voltage clamp for the indicated injections of cRNA. The whole cell currents were generated by holding the potential at −80 mV and then jumping to voltages ranging from −140 mV to +60 mV in 10 mV increments (1:1) or to +100 mV (WT and Blank). (***B***) Plots of relative conductance versus the voltage of the steps following: injection of: a 1:1 mixture of mutant and WT cRNA (red diamonds); injection of WT cRNA alone (black circles); or injection of carrier only (purple triangles). The high conductance at negative potentials would act to drive the membrane potential to about −80 mV, the equilibrium potential of K^+^. Channel activation is left shifted in the whole cell recordings in this figure compared to the ~0 Ca^2+^ macro patch recordings in Fig. 3, because the resting free Ca^2+^ in the oocytes of a few micromolar left shifts BK channel activation. The relative cord conductance was calculated from the plotted steady-state currents in Supplementary Fig. 2 using the step potential minus the reversal potential for the voltage driving force. Mean ± SEM, n = 4 in each case.

The negative shift in activation for currents expressed from the 1:1 injection was so pronounced that large numbers of aberrant BK channels would have been open at voltages near the resting membrane potential of −60 mV. If aberrant currents expressed similarly in neurons, the flux of K^+^ through opened BK channels would act to drive the membrane potential towards the equilibrium potential for K^+^, opposing depolarization of the cell and facilitating repolarization, which could interfere with normal neuronal function. Thus, at the level of whole cell and macro patch recording, the heterozygous G375R mutation gave a pronounced gain-of-function (GOF) phenotype because much less depolarization was required to activate BK currents. Some of the disease phenotypes associated with this mutation might be due to such GOF behavior. In contrast, Liang et al. reported a LOF for G375R because they observed no BK currents for homomeric mutant channels^25^. Their conclusion of a LOF vs. our conclusion of a GOF will be considered in detail in a later section.

To quantify the average negative voltage shift in activation for the BK currents following injection of a 1:1 mixture of G375R and WT cRNA, when compared to WT currents following injection of only WT cRNA, we recorded currents from macro patches of membrane which were excised from oocytes after channel expression. This allowed the composition of the solution at the inner membrane surface to be controlled, which was not the case for the whole cell recordings in Fig. 2. For a solution at the intracellular membrane surface containing <0.01 μM Ca^2+^ (~0 Ca^2+^), WT BK currents did not activate appreciably until the membrane potential exceeded 100 mV (black circles), whereas BK currents from the 1:1 injection of G375R mutant and WT cRNAs started to activate at very negative voltages of −80 mV (Fig. 3B). The mean voltage for half activation (*V*_h_) was 182 ± 5.4 mV (n = 6) for WT BK currents and 61.5 ± 16 mV (n = 12) for BK currents from the 1:1 injection, giving a mean left shift of −120 mV in voltage activation for the currents following a 1:1 injection when compared to WT (p <0.0001). This left shift in voltage activation was even greater in the presence of 300 μM Ca^2+^ (Supplementary Fig. 3). The 1:1 injection also decreased the voltage sensitivity of activation compared to WT, with a slope of 38.5 ± 1.8 (n = 8) mV per e-fold change compared to 23.2 ± 1.36 mV (n = 6) for WT currents (p < 0.0001).

**Fig. 3.**
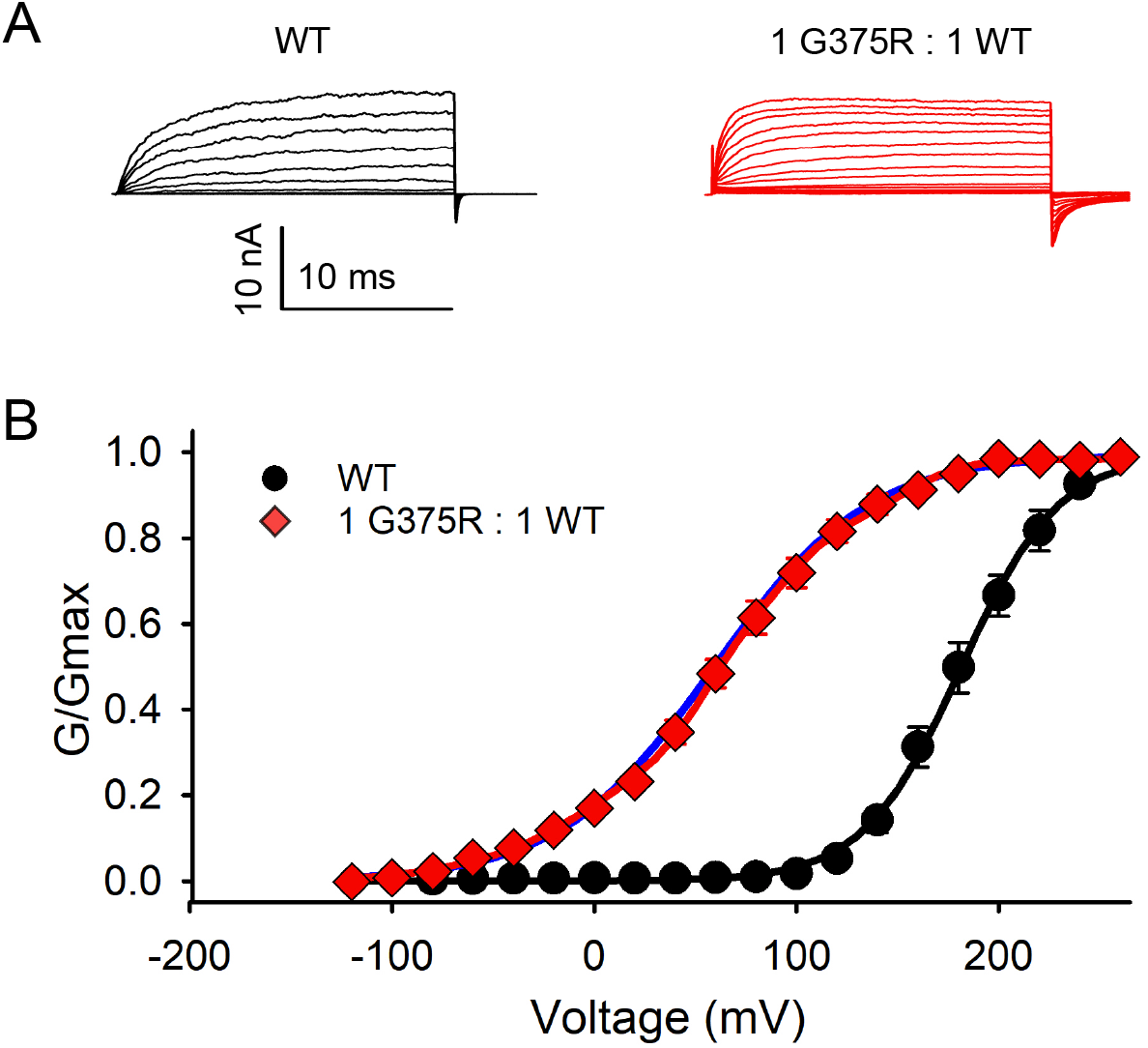
Quantifying the mean negative shift in *V*_h_ following injection of a 1:1 mixture of G375R mutant and WT cRNA when compared to injection of only WT cRNA. Data are for essentially 0 Ca^2+^ at the intracellular side of the membrane. (***A***) Currents recorded from inside-out macro patches of membrane excised from oocytes. For WT channels following injection of only WT cRNA, holding potential was −80 mV and voltage pulses were from −80 mV to 240 mV with 20 mV steps followed by a step back to −80 mV to elicit tail currents for G/Gmax measurements. For channels expressed following injection of a 1:1 mixture of G375R mutant and WT cRNA, holding potential was −160 mV and voltage pulses were from −120 to 240 mV with 20 mV steps followed by a step back to −120 mV to elicit tail current for measurement. The currents are the mean response from the many hundreds to thousands of BK channels in each excised macro patch of membrane. (***B***) G/G_max_ vs. voltage plots following the injection of WT cRNA alone (black circles, n = 6), or 1:1 injection of mutant and WT cRNA (red diamonds, n = 12). The black line through the WT data is a single Boltzmann function with *V*_h_ = 181.3 mV and slope factor *b* = 24.04 mV/e-fold change in Po at low Po. The blue line through the 1:1 data (red diamonds) is a single Boltzmann function with *V*_h_ = 59.04 mV and *b* = 38.06 mV/e-fold change. The red line through the 1:1 data is the sum of five Boltzmann functions with: 1) *V*_h_ = −75.7 mV, *b* =21.6 mV/e-fold change, fractional area = 0.0547; 2) 1.5, 21.0, 0.236; 3) 54.6, 13.8, 0.316; 4) 96.6, 14.4, 0.244; 5) 155, 14.0, 0.139. These parameters for the five summed Boltzmann functions that describe the 1:1 data are generally consistent with the *V*_h_ values for the apparent clusters of assembled channels (Fig. 5A) and theoretical percentages in Fig. 1.

The large negative shifts in voltage activation observed for whole cell recordings following a 1:1 injection of mutant and WT cRNA (Fig. 2; Supplementary Fig. 2) are thus also observed for macro patch recordings under conditions in which the intracellular solution and Ca^2+^ are controlled (Fig. 3; Supplementary Fig. 3).

Whole cell and macro patch recordings are useful to show the average response of many hundreds to thousands of BK channels, but they provide limited information about the properties of the underlying channels, unless all channels are identical, which is unlikely to be the case for a heterozygous mutation, as shown in Fig. 1. To investigate the channels underlying the negative shifts in BK currents in Figs. 2 and 3 for a 1:1 injection of mutant and WT cRNA to mimic a mutation heterozygous with the WT allele, we first consider the possible channel types and frequencies of expression that might be expected for a heterozygous mutation. We then record from individual channels expressed following a 1:1 injection of mutant and WT cRNA to see if the expectations are met.

### BK channels of five different potential stoichiometries and six different subunit arrangements could theoretically be expressed for a mutation heterozygous with WT

WT and mutant subunits arising from a heterozygous mutation could potentially assemble into tetrameric BK channels with five different stoichiometries and six different subunit arrangements, each with potentially different properties (Fig. 1)^29–33^. If it is assumed that there is equal production of mutant and WT subunits, that subunits assemble randomly to form tetrameric channels, and that each assembled channel has the same probability of reaching the surface membrane, then the channels expressed for a heterozygous mutation would consist of 6% WT channels, 25% comprised of 1 mutant and 3 WT subunits, 38% comprised of 2 mutant and 2 WT subunits, 25% comprised of 3 mutant and 1 WT subunit, and 6% homomeric mutant channels (Fig. 1). On this basis, 94%, of the expressed channels could be pathogenic, comprised of 88% hybrid (heteromeric) channels of mixed subunits and 6% homomeric mutant channels, with only 6% WT channels (Fig. 1).

### Assembled channels

The term assembled channels will be used to refer to those BK channels that are assembled and expressed following injection of a 1:1 mixture of G375R mutant and WT cRNA into *Xenopus* oocytes, to mimic a heterozygous mutation. Assembled channels could theoretically include five types of channels with different stoichiometries and six types of channels if subunit arrangement is considered (Fig. 1). An individual assembled channel could then be any one of the six channel types. It is the combined activity of the six potential types of assembled channels that would determine the BK currents for a heterozygous mutation.

### Assembled channels display an abnormally wide range of *V*_h_

To explore if multiple channel types contribute to the negative shift in *V*_h_ resulting from a G375R BK subunit mutation heterozygous with WT (Figs. 2, 3, Supplementary Figs. 2, 3), we used the unique ability of the patch clamp technique^31,35–37^ to isolate and record from individual assembled channels expressed following a 1:1 injection of mutant and WT cRNA into *Xenopus* oocytes. For each of the 33 individual assembled channels studied, single-channel currents were recorded over a range of voltages (Fig. 4*A*) to obtain plots of open probability (Po) vs. voltage (Fig. 4*B*, red curves). The Po-V plots for the 33 individual assembled channels had a voltage for half activation (*V*_h_) that ranged from −152 mV to +151 mV, for a span of 303 mV (Fig. 4*B*, red Po-V curves). This very wide range in *V*_h_ supports the idea that there are multiple types of assembled channels with different properties arising from different subunit composition (Fig. 1).

**Fig. 4.**
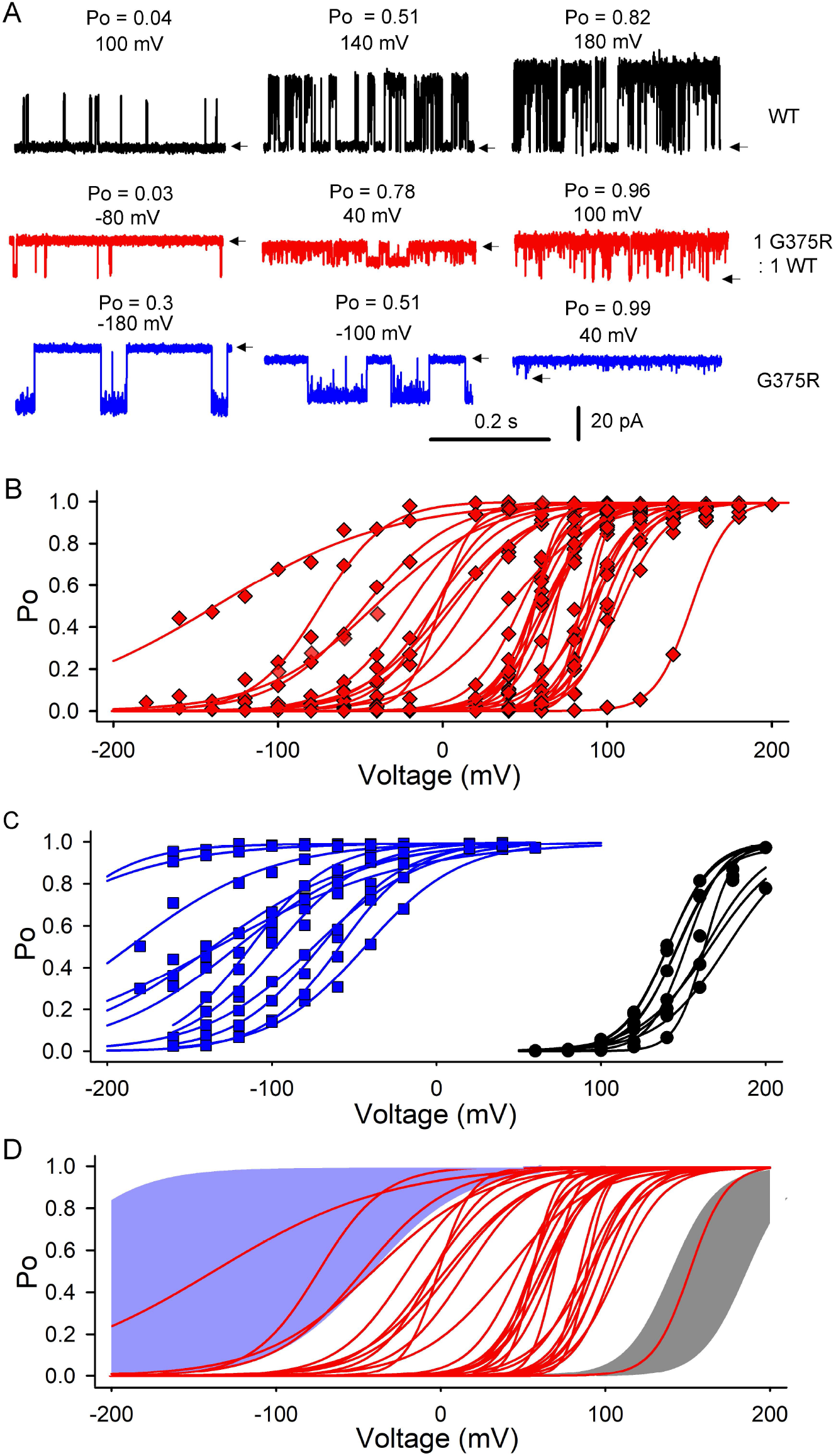
Identifying the types of assembled BK channels expressed following a 1:1 injection of G375R mutant and WT cRNA. (***A***) Representative single channel recordings from three different single BK channel molecules: WT channel (top row) following injection of WT cRNA; an assembled channel following 1:1 injection of G375R mutant and WT cRNA, (second row); and a G375R homomeric mutant channel following injection of G375R mutant cRNA (third row). Recordings are shown at three different voltages for each channel type. Depolarization activates each channel type with a markedly different *V*_h_ of 140 mV for the WT channel, 6 mV for the assembled channel, and −100 mV for the G375R homomeric mutant channel. Arrows indicate closed channel current levels. (***B***) Plots of Po vs. *V* for 33 individual assembled channels following a 1:1 injection of mutant and WT cRNA (red diamonds with Boltzmann fits). (***C***) Plots of Po vs. *V* for 12 single G375R homomeric mutant channels following injection of G375R mutant cRNA (blue squares with Boltzmann fits), and for 9 WT channels following injection of WT cRNA (black circles and Boltzmann fits). (***D***) Identifying the types of assembled channels. Blue and gray areas indicate the range of observed *V*_h_ values for G375R homomeric mutant channels and WT channels, respectively, from *C*. One of the assembled channels (rightmost red Po-*V* curve) overlaps with the WT channels (gray shading), indicating that this assembled channel is likely a WT channel with four WT subunits. Four of the assembled channels (four leftmost red Po-*V* curves) overlap with the G375R homomeric mutant channels (blue shading) indicating that these four assembled channels are likely G375R homomeric mutant channels assembled from four mutant subunits. The remaining 28 assembled channels would be hybrid channels with mixed subunits (Fig. 1), as their *V*_h_ values do not overlap with those of either homomeric mutant or WT channels.

### Assembled channels include WT, homomeric mutant, and hybrid channels

To identify the types of BK channels expressed following a 1:1 injection of mutant and WT cRNA, the Po-V curves of the 33 individual assembled channels in Fig. 4B were compared to Po-V curves obtained from individual WT and homomeric mutant channels. The WT and homomeric mutant channels were obtained by single channel recording after injection of only WT cRNA or only mutant cRNA, respectively. The WT channels had *V*_h_ values that spanned a narrow range from +140 to +176 mV (Fig. 4C, black Po-V curves), and the homomeric mutant channels had *V*_h_ values that ranged from −272 mV to −38 mV (Fig. 4C, blue Po-V curves). The homomeric mutant channels had a surprisingly wide range of *V*_h_ for channels of presumed identical composition of four mutant subunits, but all were in a far more negative voltage range than WT channels. The reason for such a wide voltage range in *V*_h_ for the homomeric mutant channels is not known, but perhaps channels with four mutant subunits can assume different conformations with markedly different activation properties, depending on mutant side-chain orientation (Supplementary Fig. 1).

To facilitate the identification of the types of assembled channels, the Po-V curves of the individual 33 assembled channels from Fig. 4B were overlaid on plots of the *V*_h_ ranges of the Po-V curves for known WT (gray shading) and homomeric mutant (blue shading) channel controls in Fig. 4D. One of the 33 assembled channels had a Po-V curve that overlapped with the known WT channel controls (Fig. 4D), suggesting that this assembled channel was WT with four WT subunits. Four of the 33 assembled channels had Po-V curves that overlapped with the known homomeric mutant channel controls (Fig. 4D), suggesting that these four assembled channels were homomeric mutant, comprised of four mutant subunits. The remaining 28 assembled channels had Po-V curves that did not overlap with the known WT or homomeric mutant channel controls (Figs. 4D), but fell in between, suggesting that these 28 assembled channels were all hybrid channels comprised of a mix of mutant and WT subunits, because hybrid channels were the only types of channels remaining (Fig. 1). In this study, assembled channels with *V*_h_ values in the ranges of WT, hybrid, and homomeric mutant channels will be referred to as WT, hybrid, and homomeric mutant channels, respectively, with the understanding that these classifications are based on *V*_h_ values.

For the sampled group of 33 assembled channels, ~3% (1/33) were WT, ~85% (28/33) were hybrid, and ~12% (4/33) were homomeric mutant (Fig. 4B-D). Fig. 1 predicts ~6% WT, ~88% hybrid, and ~6% homomeric mutant channels. Thus, for the G375R mutation heterozygous with WT, both experimental and theoretical considerations suggest that most (85-88%) of the expressed assembled channels will be hybrid, with much smaller fractions of homomeric mutant and WT channels. All hybrid and homomeric mutant channels displayed negative shifts in activation compared to WT, with the greatest negative shifts for the homomeric mutant channels. Consequently, both theoretical and experimental considerations suggest that 94-97% of the BK channels arising from a G375R mutation heterozygous with WT would display aberrant negative shifts in activation, even though only 50% of the subunits synthesized in a cell would be mutant.

### Three functional types of hybrid assembled channels

Assembled channels were found to be comprised of 85% hybrid channels, 3% WT, and 12% homomeric mutant (Fig. 4). The 85% hybrid channels themselves could consist of three types, with one, two, or three mutant subunits per channel replacing an equal number of WT subunits (Fig. 1). To explore this possibility, a histogram of the *V*_h_ values of the 33 assembled channels from Fig. 4B was plotted in Fig. 5A as red bars. Histograms of known WT channel controls (black bars) and homomeric mutant channel controls (blue bars) from Fig. 4C are also plotted. The subunit compositions of the homomeric mutant and WT channel controls are known and placed above these two types of channels in Fig. 5A. As expected from Fig. 4D, four of the assembled channels had *V*_h_ values that overlapped with those of homomeric mutant channels, suggesting that they were homomeric mutant channels; one assembled channel overlapped with WT, suggesting that it was a WT channel; and the remaining 28 assembled channels had *V*_h_ values falling in between those of homomeric mutant and WT channels, identifying them as hybrid channels (Fig. 1).

**Fig. 5.**
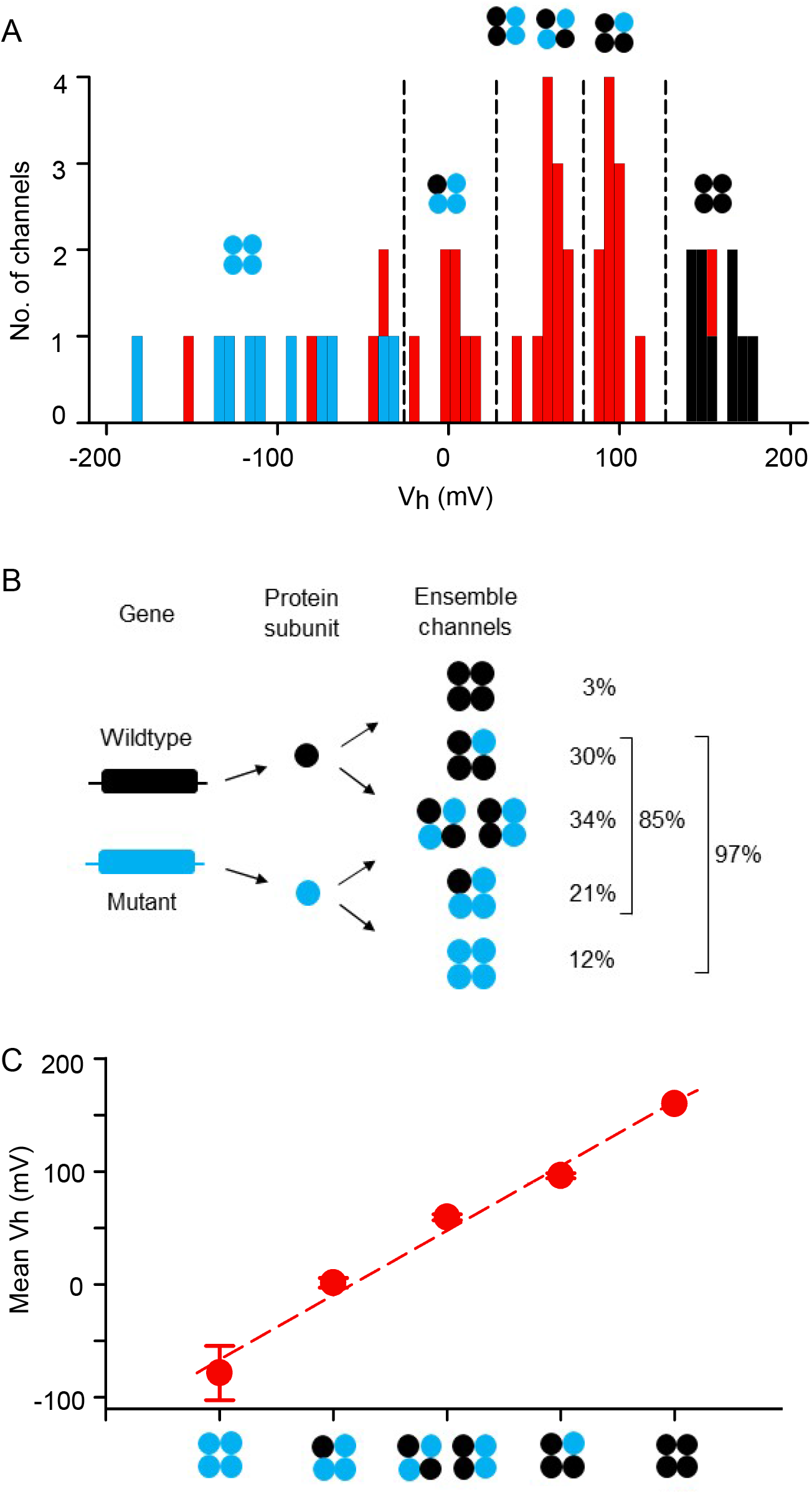
Five different types of functional BK channels are observed following injection of a 1:1 mixture of mutant and WT cRNA into *Xenopus* oocytes. *(**A**)* Histograms of *V*_h_ values of single channels from Fig. 4. WT channels expressed after injection of only WT cRNA are indicated by black bars, and G375R homomeric mutant channels expressed after injection of only G375R cRNA are indicated by blue bars. Assembled channels expressed after a 1:1 injection of mutant and WT cRNA are indicated by red bars. The red bar in the cluster of black bars is an assembled channel comprised of four WT subunits. The four red bars interspaced between the blue bars are assembled channels comprised of four mutant subunits. The 28 red bars with *V*_h_ values in between the blue bars and black bars are hybrid assembled channels of mixed subunits. The hybrid channels appear to cluster into three groups with mean *V*_h_ values of 1.4, 59 and 96 mV. Hypothesized subunit compositions of the three apparent clusters of hybrid channels are indicated, where black indicates WT subunits and blue indicates G375R mutant subunits. (***B***) The experimentally observed frequencies of the five types of assembled channels are listed for comparison to the theoretical frequencies in Fig. 1. (***C***) *V*_h_ for the five types of assembled channels is approximated by a linear incremental model where each subunit contributes an increment of *V*_h_,

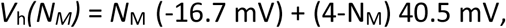

where *N*_M_ is the number of G375R mutant subunits per channel, *V*_h_*(N_M_)* is *V*_h_ as a function of *N_M_*, and −16.7 mV and 40.5 mV are the increments of *V*_h_ added by each mutant and WT subunit, respectively. Filled red circles are the experimentally observed mean ± SEM values of *V*_h_ for the five types of assembled channels from part *A*, and the dashed line indicates the predicted values. Predictions are for 0 intracellular Ca^2+^. Mean ± SEM. Histograms of single-channel kinetic parameters have been used previously to investigate the number of β2 regulatory subunits per BK channel^50^.

The *V*_h_ values of the 28 hybrid channels fell into three apparent clusters with mean values of about 1.4, 59 and 96 mV (Fig. 5A). Three clusters of *V*_h_ for hybrid channels would be expected from Fig. 1 if each mutant subunit replacing a WT subunit added an increment of negative shift in *V*_h_, as suggested by Fig. 4 and in Fig. 6 in a later section. Accordingly, as a working hypothesis, the different subunit combinations for hybrid channels from Fig. 1 have been placed above the three clusters of hybrid channels in Fig. 5A to obtain a stepwise negative shift in *V*_h_ for each additional mutant subunit replacing a WT subunit. Because only three clusters of hybrid channels were observed, instead of four, it has been assumed that the hybrid channels with two mutant and two WT subunits, but of different subunit arrangement, contribute to the same cluster. Fig. 5A then suggests five different types of functional assembled channels: homomeric mutant, three types of hybrid channels, and WT channels, as indicated. The percentages of expression of the five functional types of assembled channels were then calculated from Fig. 5A and presented in Fig. 5B. Simulation of 1 million groups of 33 assembled channels, assuming equal production of mutant and WT subunits followed by random assembly into tetrameric channels, indicated that the experimentally observed percentages in Fig. 5B were not significantly different from the predictions of Fig. 1 (Supplementary Fig. 4).

**Fig. 6.**
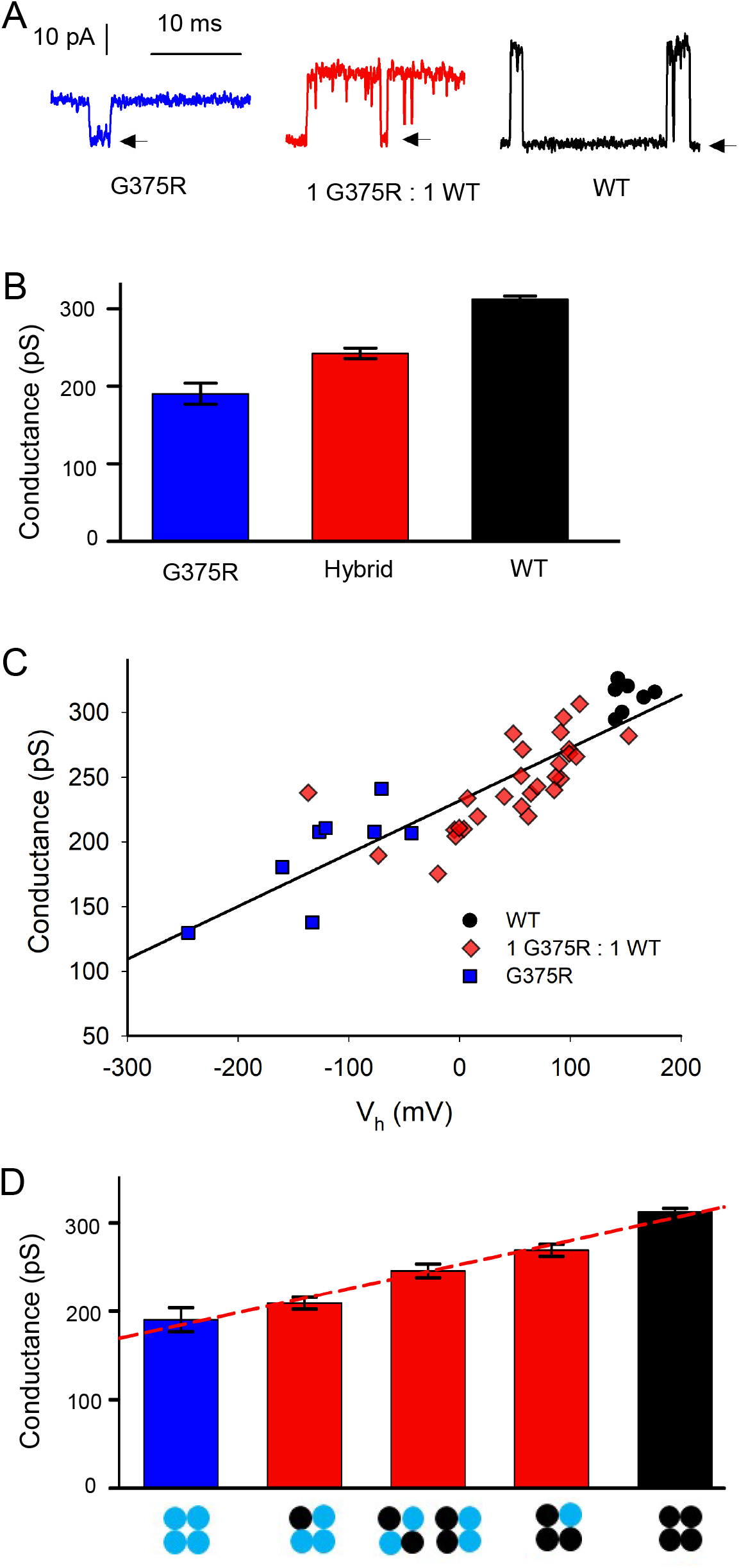
Each replacement of a WT subunit with a G375R mutant subunit in a BK channel acts to both left shift *V*_h_ to more negative potentials and decrease single channel conductance. (***A***) Single-channel currents recorded from single channels at +100 mV after injecting the indicated cRNA. (***B***) Single-channel conductance at +100 mV decreases as the number of mutant subunits increase. Mean single-channel conductance for WT channels was 312 ± 4 pS (n = 7), decreasing to 245 ± 6 pS for hybrid channels (n = 25, p < 0.0001), and further decreasing to 190 ± 14 pS for G375R homomeric mutant channels (n = 8, p = 0.0003). Mean ± SEM. WT and G375R homomeric mutant channels were those channels expressed after injecting only WT cRNA or only G375R mutant cRNA, respectively. Hybrid channels were identified as in Fig. 4D. (***C***) Plot of single channel conductance at +100 mV vs. *V*_h_ for BK channels expressed following injection of the indicated cRNA. The linear regression line plots single channel conductance *g* vs. *V*_h_, where: *g* = *g*(0) + *aV*, where *g*(0) = 231.9 ± 3.9 pA and *a* = 0.408 ± 0.0375 pS/mV (p <0.0001 for slope significantly different from 0. R = 0.86). (***D***) Single channel conductance for the five types of assembled channels is predicted by a linear incremental model, where each subunit contributes an increment of conductance,

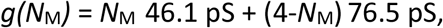

where *N*_M_ is the number of G375R mutant subunits per channel, *g(N*_M_) is single channel conductance as a function of *N_M_*, and 46.1 pS and 76.5 pS are the increments of *g* added by each mutant and WT subunit, respectively. The homomeric mutant and WT data are from part *B*, and the three red hybrid conductance bars plot *g* from the three groups of hybrid channels in Fig. 5A. The red dashed line indicates the predicted values of *g.* Mean ± SEM.

Whereas the observation of three apparent clusters of *V*_h_ values for the hybrid assembled channels (Fig. 5) is consistent with theoretical predictions (Fig. 1), peaks can occur by chance alone in histograms of binned data of limited sample size^38^. Consequently, additional experiments would be needed to determine whether *V*_h_ values for hybrid channels can consistently be resolved into three peaks, but an observation of such distinct peaks is not required for support of Fig. 1, as variability in *V*_h_ among channels of the same type^39^ might be sufficient to obscure distinct peaks.

### Linear Incremental model for the contributions of mutant and WT subunits to *V*_h_

WT BK channels comprised of four WT subunits had a mean *V*_h_ of about 160 mV, and homomeric mutant BK channels comprised of four mutant subunits had a mean *V*_h_ of about −80 mV (Figs. 4A, C; Fig. 5A). As all four subunits contribute to the gating of BK channels^4,5,31^, these observations suggest that each WT subunit contributes an increment of positive shift to *V*_h_ and each mutant subunit contributes an increment of negative shift. In support of this hypothesis, a linear incremental model in which each WT subunit in a channel added +40.5 mV to *V*_h_ and each mutant subunit added −16.7 mV, provided an approximate description of the observed *V*_h_’s for the five types of assembled channels (Fig. 5C dashed line; model in figure legend). The net effect of replacing a WT subunit with a mutant subunit was a −57.2 mV shift in *V*_h_ to account for the loss of the positive shift contributed by the removed WT subunit and the addition of the negative shift contributed by the added mutant subunit.

### Dual action of G375R subunits on *V*_h_ and single channel conductance *g*

In addition to the negative shift in activation induced by replacing WT subunits with G375R mutant subunits (Figs. 2–5), replacing WT subunits with mutant subunits also decreased single-channel conductance (Fig. 6A, B). The mean single-channel conductance of WT channels was 312 ± 4 pS. This decreased to 245 ± 6 pS for hybrid channels, and further decreased to 190 ± 14 pS for homomeric mutant channels. These decreases were significant (Fig. 6 legend). Hence, single-channel conductance decreased as mutant subunits replaced WT subunits. The decreased single-channel conductance may arise from the larger volume and positive charge of the mutant arginine side chains replacing the single hydrogen atom of the glycine side chains on the four S6 segments lining the conductance pathway of the BK channel (Supplementary Fig. 1). The added volume of the sidechains could decrease the volume of the inner vestibule and the added positive charge may act to repel K^+^ from the inner cavity. Both actions can reduce single channel conductance in BK channels^40–42^.

To examine the relationship between single-channel conductance and *V*_h_, the single-channel conductance for each channel was plotted against *V*_h_ for the same channel in Fig. 6C for the indicated channel types. A linear relationship was observed (Fig. 6C; R = 0.86, P < 0.0001). When taken together, the data in Figs. 6B and 6C are consistent with the idea that replacing a WT subunit with a mutant subunit adds both an increment of negative voltage shift to *V*_h_ and a step decrease in single-channel conductance. Thus, at the single-channel level, the heterozygous G375R mutation acts simultaneously as a GOF mutation to shift voltage activation to more negative voltages (Figs. 4, 5), and as a loss-of-function (LOF) mutation to decrease single-channel conductance (Fig. 6). At the whole cell and macro patch level, the increase in currents from the GOF negative shift in activation would dominate the LOF reduction in single-channel conductance, producing large left shifted currents (Figs. 2, 3; Supplementary Figs. 2 and 3). The LOF in conductance would act to decrease the consequences of the negative shift in activation.

### Linear Incremental model for the contributions of mutant and WT subunits to single-channel conductance

A linear incremental model in which each WT subunit contributes 76.5 pS to single channel conductance and each mutant subunit contributes 46.1 pS could approximate the single channel conductance for the five types of assembled channels (Fig. 6D dashed line; model in figure legend).

### WT, hybrid, and G375R homomeric mutant channels are expressed in the HEK293 cell expression system

Liang et al.^25^ reported that no potassium currents were recorded from excised macro patches following transfection of HEK293T cells with plasmids coding for G375R mutant BK subunits. In contrast, we found that functional single homomeric mutant channels were readily expressed following injection of *Xenopus* oocytes with cRNA coding for mutant BK G375R subunits (Fig. 4A, C). To examine if these differences in channel expression were due to differences in expression systems, we transfected HEK293 cells with WT cDNA to generate WT channels, with mutant G375R cDNA to generate homomeric mutant channels, and with a 1:1 mixture of WT and mutant cDNA to mimic the generation of channels that would normally arise from a heterozygous mutation. We found that the transfected HEK293 cells displayed robust whole-cell BK currents for each of the three different types of transfection (Fig. 7A). The G-V plots for transfection with WT cDNA or 1:1 transfection with WT and mutant cDNA were very similar to those we obtained using the *Xenopus* oocyte expression system (compare Fig. 7 with Fig. 3). We also found that whole cell G375R homomeric mutant currents from HEK293 cells had *V*_h_ values within the broad range of *V*_h_ values of single-channel recordings from patches of membrane excised from *Xenopus* oocytes (Fig. 4).

**Fig. 7.**
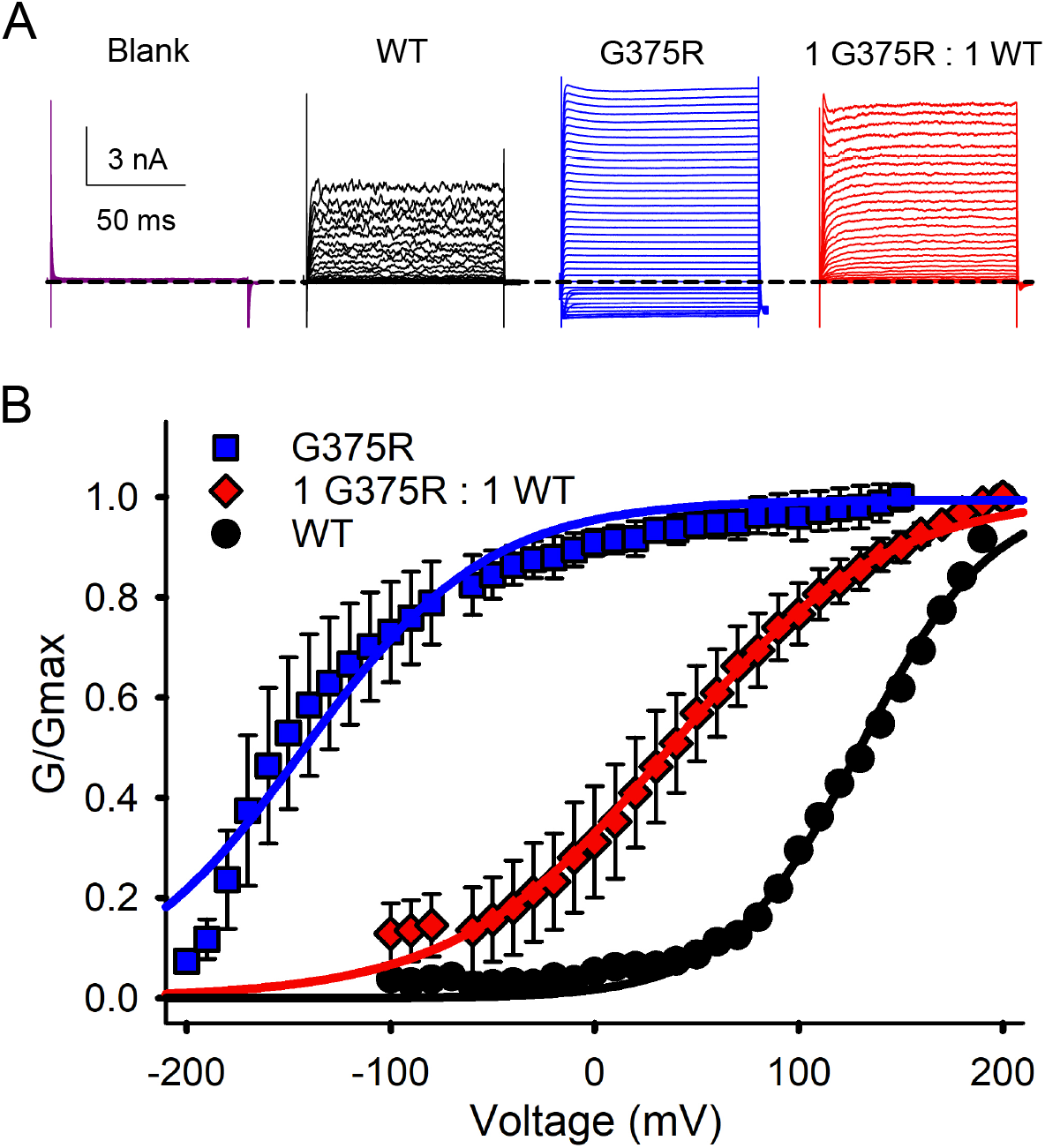
G375R homomeric mutant channels, assembled channels, and WT BK channels are expressed in HEK293 cells. (***A***) Whole cell currents recorded from HEK293 cells following transfection with the indicated types of cDNA. The whole cell currents were generated by holding at −60 mV, with a pre-step to −100 mV, followed by steps from −200 or −100 to +200 mV in 10 mV increments followed by a post-step to −120 mV. The dashed line indicates the level of 0 current. Evoked currents were not observed in HEK293 cells that were not transfected. (***B***) Plots of normalized conductance versus the voltage of the activating steps following transfections with G375R mutant cDNA (blue squares), a 1:1 mixture of mutant and WT cDNA (red diamonds) to express assembled channels, or WT cDNA (black circles). Unlike single channel recordings (Fig. 4), whole cell recordings reflect the average response from many hundreds to thousands of channels. There were large negative shifts in *V*_h_ for G/Gmax curves determined from G375R homomeric mutant channels and from assembled channels, when compared to WT channels. The mean voltage for half activation (*V*_h_) was 130.3 ± 4.4 mV (n = 4) for WT BK currents, 37.5 ± 2.7 mV (n = 5) for BK currents from the 1:1 injection, and −142.5 ± 3.6 mV (n = 3) for the G375R homomeric mutant channels. The 1:1 transfection also decreased the voltage sensitivity of activation compared to WT, with a slope of 52.0 ± 2.0 mV (n = 5) per e-fold change for current after a 1:1 transfection, compared to 32.3 ± 2.5 mV (n = 4) for WT currents. The G375R homomeric mutant channels had a slope of 45.0 ± 4.0 mV. Mean ± SEM.

Hence, in both *Xenopus* oocytes and HEK293 expression systems we observed functional G375R homomeric mutant currents that displayed a marked negative shift in activation consistent with a GOF mutation. This contrasts with the observations of Liang et al.^25^ who did not observe potassium currents for G375R transfection of HEK293T cells. They reported G375R as a LOF mutation. The reason for the difference in observations is not known, but we found that the viability of *Xenopus* oocytes injected with G375R cRNA was reduced (see Methods).

### Whole cell and macro-patch currents can obscure mechanism

The narrow error bars for the macro currents recorded from whole oocytes and excised macro patches (Figs. 2 and 3; Supplemental Figs. 2 and 3) following injection of WT cRNA or a 1:1 injection of mutant and WT cRNA, indicated that the mean responses recorded from hundreds to thousands of channels were repeatable. Narrow error bars for mean responses do not necessarily indicate that the individual channels underlying the mean response have near identical properties, as this was not the case for the macro currents following 1:1 injections of mutant and WT cRNA, where individual assembled channels for 1:1 injections had a very wide range of *V*_h_ that spanned over 300 mV (Fig. 4B). In spite of this wide range for individual channels, the mean G-V data following the 1:1 injections were reasonably well described by a single Boltzmann function (Fig. 3B, blue line through red diamonds), but were better described (red line) by assuming five underlying channel types (Fig. 1) as described in the legend to Fig. 3.

## Discussion

The heterozygous G375R BK channel variant has been associated with a devastating human phenotype that includes malformation syndrome and severe neurological and developmental disorders^25^. This variant has only appeared in the human population when heterozygous with WT, perhaps because a homozygous G375R variant may not permit viability because of the extreme GOF phenotype we observed for homomeric mutant channels. To gain insight into the pathogenicity of this variant, currents from whole cells, macro patches, and single channels were recorded from BK channels expressed following a 1:1 injection of G375R mutant and WT cRNA to mimic a G375R mutation heterozygous with WT.

Recordings from whole cells and macro patches, both of which contain many hundreds to thousands of channels, indicated that the voltage required for half activation, *V*_h_, of the current following a 1:1 injection, was left shifted to more negative potentials by about −120 mV compared to WT currents (Figs. 2 and 3; Supplementary Figs. 2 and 3), indicating a pronounced net GOF phenotype on channel activation. The aberrant BK channels underlying the negative shifts in activation would lead to a much greater fraction of BK channels being open at negative membrane potentials, including at potentials near the resting potential (Figs. 2, 3 and Supplementary Figs. 2, 3), which would oppose cellular depolarization, altering cellular function.

To explore the underlying molecular basis for the aberrant current activation associated with the heterozygous G375R variant, we used high resolution single channel recording to isolate and characterize the BK channels, termed assembled channels, that are assembled and expressed following injection of a 1:1 mixture of G375R mutant and WT cRNA. Theoretical considerations based on equal production and random assembly of mutant and WT subunits suggest there could be multiple types of assembled channels with different subunit combinations^29,30,32,33^ (Fig. 1) Consistent with this possibility, our analysis suggested that five different types of functional BK channels were expressed: 3% were WT, 12% were homomeric mutant, and 85% were three different types of hybrid channels of mixed subunits (Fig. 4, Fig. 5A, B). The frequencies of expression of these five types of functional assembled channels were not significantly different (Supplementary Fig. 4) from the theoretical predictions of Fig. 1 based on equal production and random assembly of subunits. This suggests that the processes involved in subunit production, assembly, and expression do not distinguish significantly between the mutant and WT alleles or subunits, so that phenotypic differences arise at the level of individual channels.

The three types of hybrid channels comprising 85% of the expressed assembled channels, had properties falling between those of mutant and WT channels, that varied with their apparent subunit composition (Figs. 4–6). Ninety-seven percent of the assembled channels, all except for the 3% WT, displayed both GOF negative shifts in activation and LOF decreases in single-channel conductance (Figs. 4–6). Hence, most of the expressed channels for the heterozygous G375R mutation displayed aberrant properties. The values of *V*_h_ and single channel conductance for each of the five types of functional assembled channels could be predicted with a linear incremental model in which each mutant and WT subunit in each of the five types of functional assembled channels acted relatively independently to contribute increments of both *V*_h_ and single-channel conductance to the molecular phenotype of the channel (Figs. 5C, 6D).

A potential mechanism to produce the five functional types of assembled channels for a heterozygous G375R BK channel mutation, then, is equal production and random assembly of mutant and WT subunits into channels of five different subunit compositions, where the *V*_h_ and single-channel conductance of each channel type are determined by independent contributions from each of the four subunits in a channel. Each WT subunit adds increments of 40.5 mV to *V*_h_ and 76.5 pS to single channel conductance, and each mutant subunit adds increments of −16.7 mV to *V*_h_ and 46.1 pS to single channel conductance.

To our knowledge, this is the first time that the types of functional channels expressed for a heterozygous mutation, their specific phenotypes, frequencies of expression, and contributions of individual subunits have been revealed in such molecular detail (Figs. 4–6). With this molecular information in hand, it is now possible to classify the heterozygous G375R mutation with regards to inheritance of genetic disease, and also the individual molecular phenotypes of the five types of expressed functional channels.

For classification with regard to genetic disease, we suggest that the heterozygous G375R mutation be labeled as an assembly-mediated dominant-positive mutation, following the terminology of Backwell and Marsh^33^. Assembly-mediated to indicate that mutant and WT subunits assemble into multiple types of functional channels for the tetrameric BK channel, dominant because the inclusion of only a single copy of the mutant *KCNMA1* gene leads to disease^25^, and positive to indicate an aberrant GOF in the activation of BK currents (Figs. 2, 3, 7). The net result of this assembly-mediated dominant-positive mutation in the tetrameric BK channels is that a disproportionate fraction (94-97%) of the expressed channels display aberrant activation due to the random assembly of mutant and WT subunits into functional channels (Figs. 1, 4, 5). A dominant-positive GOF phenotype has been described previously for the heterozygous G88R mutation of the TASK-4 K^+^ channel in the heart, based on a study using whole cell currents^43^.

A dominant-positive effect is an analogy^33^ to the well-known dominant-negative effect observed for some types of protein complexes and channelopathies^33,44–49^ Both can cause disease: dominant positive from a GOF and dominant-negative from complete or partial LOF, and both have a disproportionate fraction of the channels affected.

Whereas an assembly-mediated dominant-positive classification of the heterozygous G375R mutation is useful to suggest a basis of genetic disease at the level of whole cell and macro patch currents through BK channels, it describes only part of the molecular phenotype. Consequently, to characterize the molecular phenotype we suggest that random assembly of mutant and WT subunits for the heterozygous G375R mutation produces five functional types of assembled channels: WT, mutant, and three types of hybrid channels of mixed subunit composition. All channel types except WT simultaneously display a GOF in activation and a partial LOF in single-channel conductance.

The genetic dominance of the various channel types expressed for a heterozygous mutation is mixed and complex. Codominance was observed for the WT and homomeric mutant channels and partial dominance was observed for the three types of hybrid channels. Support for codominance is that the mutant and WT channels expressed when mimicking a heterozygous mutation had the same molecular phenotypes as homomeric mutant and WT channels expressed after injection of only mutant or WT cRNA, respectively (Figs. 4 and 5). Support for partial dominance is that the three types of hybrid channels had their own values of *V*_h_ and single channel conductance that were distinct from each other and fell in the range between those of the homomeric mutant and WT channels (Figs. 5C, 6D). The levels of partial dominance in the linear incremental model (Figs. 5C and 6D) were determined by the numbers of mutant and WT subunits per channel acting relatively independently of one another, rather than by mutant subunits altering the function of WT subunits, as in some classical descriptions of dominant-negative. It is remarkable that a single base pair substitution in one allele of a pair of alleles that encode for the alpha subunit of BK channels would result in WT channels plus four aberrant channel types, each with different molecular phenotypes that simultaneously display dominant positive and dominant negative effects, with the five types of functional channels displaying either co-dominance or one of three levels of partial dominance.

How do our observations of a relatively independent interactions of the G375R mutant and WT pore-forming subunits compare to interactions of these subunits with regulatory subunits? Wang et al.^50^ found that single BK channels comprised of four subunits could be associated with from 0 to 4 regulatory β2-subunits per channel, with each β2-subunit giving incremental changes in *V*_h_. Hence both mutated and WT pore forming subunits^36^ (Fig. 5) and the non-pore forming regulatory β2-subunits^50^ can give incremental changes in gating, but such incremental changes per subunit are not necessarily universal, so each type of subunit will need to be checked. A single BK channel comprised of four subunits can also include up to four regulatory γ1-subunits, but a single γ1-subunit per channel is sufficient to induce the full gating shift induced by γ1-subunits^51,52^.

Devising therapies for the *de novo* G375R heterozygous mutation will be challenging. Theoretical and experimental observations (Figs. 1, 4, 5) suggest that most (94 to 97%) of the channels for a mutation heterozygous with WT would be pathogenic, with only 3 to 6% of the channels WT. The pathogenic channels would consist of multiple channel types, each with large differences in activation properties and smaller differences in conductance (Figs. 4 - 6). Therapies to block or inactivate the pathogenic channels would ideally silence the multiple types of pathogenic channels while leaving any WT channels intact. Even if such selective blockers could be devised, it is unlikely that the remaining 3 to 6% of WT channels would be sufficient to restore normal cellular function. Effective therapies will likely require replacing or silencing the mutant alleles, or preventing mutant subunits from assembling with themselves and WT subunits if/when such techniques become practical in humans.

## Methods

### Constructs

Experiments were performed using the human large conductance calcium-activated potassium (BK) channel (hSlo1) KCNMA1 transcript: GenBank: U23767.1^3^ for WT, and a G375R mutation (VFFIL**G**GLAMF to VFF IL**R**GLAMF) was constructed to match the G375R *de novo* variant described by Liang et al.^25^. G375R is at the same position as G310 in Tao and MacKinnon^15^, which they called the gating hinge residue. The numbering differs because Tao and MacKinnon^15^ used an alternative transcript missing the first 65 amino acids compared to U23767.1. Both our hSlo1 WT cRNA plasmid and WT mammalian transfection plasmid contain the same hSlo1 insert and are described in McCobb et. al.^3^. Overlap extension PCR cloning was used to generate the G375R mutation, which was verified by sequencing. The new construct was linearized downstream of the end of coding and transcribed with T3 using Invitrogen’s T3 mMessage mMachine kit to make cRNA for injection into *Xenopus* oocytes. To mimic the effects of a heterozygous mutation, G375R mutant and WT cRNA were mixed in a 1:1 ratio by weight before injection into oocytes. For expression in HEK293 cells, two mammalian transfection plasmids were created. DNA fragments containing the complete channel cDNAs of either the WT or G375R mutant and two unique restriction sites flanking it were amplified by PCR and ligated into mammalian transfection plasmids, which were verified by sequencing.

### Expression of BK channels in oocytes for whole cell recording

Defolliculated *Xenopus* oocytes were injected with 0.5-150 ng of cRNA using a Nanoject II (Drummond Scientific) and incubated at 18 °C for 2-5 days before recording. Incubation was in ND96 complete medium consisting of (mM) 96 NaCl, 2 KCl, 1.8 CaCl_2_, 5 MgCl_2,_ and 5 HEPES adjusted to pH 7.5, supplemented with 2.5 mM sodium pyruvate and 100 μg/ml each penicillin and streptomycin. The two-microelectrode whole cell voltage-clamp recordings from oocytes were obtained in ND96 medium with 1 mM added 4,4′-diisothiocyanatostilbene-2,2′-disulphonic acid disodium salt hydrate (DIDS) to block the endogenous chloride conductance. Currents were obtained with an Oocyte Clamp OC-725C amplifier (Warner Instrument Corp.) Recordings were low-pass filtered at 1 kHz and digitized at 10 kHz. Electrodes were made with borosilicate glass capillaries (World Precision Instruments) pulled with a Sutter Instrument Co. P-87 pipette puller and filled with 3 M KCl.

### Macro patch and single-channel recordings from BK channels expressed in *Xenopus* oocytes

Oocytes were injected with 0.1-18 ng of cRNA and incubated at 18°C for 2-5 days in Barth’s Solution (mM): 88 NaCl, 1 KCl, 2.4 NaHC0_3_, 0.33 Ca(NO_3_)_2_, 0.41 CaCl_2_, 0.82 MgSO_4_, 15 mM HEPES, pH 7.6, plus 12 μM tetracycline. Macro and single channel currents were recorded from inside-out patches of membrane^35^ excised from oocytes at room temperature (21°C to 24°C). pClamp 9.0 software (Molecular Devices) was used to drive an Axopatch 200B amplifier to collect the currents. For macro patch current recordings, borosilicate pipettes with 0.5–2 MΩ resistance were used. The macro currents were filtered at 10 KHz and sampled at 100 kHz. A minus P/4 protocol was used to remove capacitive transients and leak currents. For single-channel recordings, borosilicate pipettes with 8-12 MΩ resistance were used. The single-channel currents were filtered at 5 KHz and sampled at 200 KHz. The pipette (external) solution contained (mM): 160 KCl, 2 MgCl_2,_ 5 TES buffer, pH 7.0. The internal membrane surface of the excised patches was perfused by two different solutions. The designated 0 Ca^2+^ solution had a free Ca^2+^ <0.01 μM and contained (mM): 160 KCl, 1 EGTA, 1 HEDTA, and 10 HEPES, pH 7.0. The solution with 300 μM internal Ca^2+^ contained (mM): 160 KCl, 0.3 CaCl_2_, 10 HEPES, pH 7.0. Procedures to obtain oocytes from *Xenopus* laevis were approved by the University of Miami Animal Care and Use Committee. Macro patch and single channel recordings were analyzed with Clampfit 10.7 software (Molecular Devices) and SigmaPlot 12.

For injection of only G375R cRNA into *Xenopus* oocytes we found that it was difficult to get giga-ohm seals of sufficient quality for macro patch recordings using the lower resistance electrodes required for such recordings. Hence, macro patch currents are not presented for injection of only G375R cRNA into oocytes. However, it was still possible to obtain high quality giga-ohm seals for single-channel recording using the higher resistance pipettes required for such recordings (Fig. 4). We also found that the viability of the oocytes was greatly reduced following injection of only G375R cRNA, with the oocytes often starting to die by the second or third day after injection, rather than after a week or more, perhaps because a large fraction of the G375R homomeric mutant channels would be expected to be open at resting membrane potentials (Figs. 4, 7). These viability problems were not observed for injection of 1:1 mixtures of G375R mutant and WT cRNA, which gave less pronounced negative shifts in activation or for injections of only WT cRNA (Figs. 3, 4, 7),

The open probability (Po) of each single channel analyzed in detail was determined for a range of voltages with Clampfit 10.7 using 50% threshold analysis to measure open and closed interval durations. Po at each voltage was calculated by dividing the total open time by the sum of the open and closed times. The durations of recordings to estimate Po ranged from ~ 5 s to ~3 min with the time increasing as the Po decreased. For macro patch current recordings, relative conductance was determined from macroscopic tail current amplitudes using the voltage protocols indicated in the figure legends. G/G_max_ vs. V (G-V) plots for macro current recordings and Po vs. V (Po-V) plots for single channels were fitted with a Boltzmann function to estimate *V*_h_, the voltage required for half maximal activation, and *b*, a measure of voltage sensitivity, using

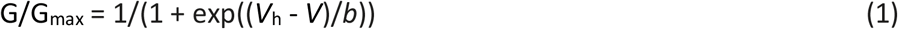

where G/G_max_ is the ratio of conductance to maximum conductance and *b* is the slope factor which gives a measure of voltage sensitivity, where *b* indicates the change in millivolts required to increase G/G_max_ (or Po for single channel recording) *e*-fold at very low G/G_max_ (or Po). Note that an increase in *b* indicates a decrease in slope and voltage sensitivity.

Single-channel conductance was determined at 100 mV by setting horizontal cursor lines by eye to the open and closed current levels of single-channel recordings.

### HEK293 cell culture

As described previously^53^, Human Embryonic Kidney HEK293 cells (ATCC CRL-1573) (ATCC, Manassas, VA) were cultured in Gibco Dulbecco’s modified Eagle’s medium (DMEM) with 10% fetal bovine serum, 100 units ml^−1^ penicillin and 100 μg ml^−1^ streptomycin (ThermoFisher Scientific, Waltham, MA) and incubated at 37°C with 5% CO_2_. These cells were grown and passaged twice a week in T25 flasks (MidSci, St. Louis, MO).

### Expression of BK channels in HEK293 cells

HEK293 cells were plated at a density of ~400,000 cells per 40 mm petri dish (TPP catalog #93040) a day before transfection. For transfection, plasmid DNA containing cDNA encoding for BK WT and/or BK G375R (mutant) subunits was added to cell layers that were 70-90% confluent using Lipofectamine 2000 transfection reagent (catalog #11668-027, ThermoFisher Scientific, Waltham, MA) following manufacturer’s instructions. 2 μg cDNA of WT, mutant, or a mix of WT and mutant (1:1 ratio by weight) was transfected per 40 mm dish. As a marker for transfection, cells were co-transfected with pmaxGFP a CMV plasmid expressing green fluorescent protein (Amaxa Biosystems), at 0.2 μg per dish. After transfection, the cells were incubated at 37°C with 5% CO_2_ for 2-4 days until recording. The recordings were then performed at room temperature (22°C). Fluorescent cells were used to study the function of the ion channels that the HEK293 cells were transfected with; non-fluorescing cells were used as a non-transfected controls^54^.

### Whole cell recording from HEK293 cells

Whole cell recordings were obtained from HEK293 cells using an Axopatch 200B amplifier (Molecular Devices, San Jose, CA). Recordings were filtered at 5 kHz with the amplifier internal filter and digitized at 50 kHz using a Digidata 1550B digitizer (Molecular Devices, San Jose, CA). The recording pipette was filled with (in mM): 140 KMES, 5 EGTA, 10 HEPES, pH 7.4 with KOH. Bath solutions contained (in mM): 135 NaMES, 5 KMES, 1 MgCl_2_, 10 HEPES, pH 7.4 with NaOH.

### Calculating the percentages of the five different stoichiometries of BK channels that can be expressed when mimicking a heterozygous mutation

As shown in Fig. 1, a mutation heterozygous with WT can potentially produce five different types of channels, defined by their stoichiometry. The binomial equation was used to calculate the expected percentages of different combinations of mutant and WT subunits for tetrameric BK channels^29,30^. For the calculations in Fig. 1 this approach assumes that WT and mutant alleles produce equal numbers of mutant and WT subunits that assemble randomly into tetrameric channels of different stoichiometries, labelled assembled channels in Fig. 1, and that each assembled channel has the same probability of being expressed, independent of subunit composition, such that

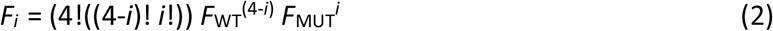

where *Fi* is the fraction of channel types with *i* mutant subunits, *F*_WT_ is the fraction of WT cRNA injected, and *F*_MUT_ is the fraction of mutant cRNA injected, where *F*_WT_ + *F*_MUT_ = 1. For BK channels, *i* ranges from 0 to 4, giving five channel types based on subunit composition. Setting both *F*_WT_ and *F*_MUT_ to 0.5 gives the fractions of the various channel types for a heterozygous mutation.

### Simulating the discrete probability distributions of the number of BK channels of each stoichiometry expected for equal production and random assembly of mutant and WT subunits

The simulated discrete probability distributions were used to determine if the experimentally observed number of channels of each stoichiometry differed significantly from the number expected for random assembly of mutant and WT subunits. Discrete probability distributions for the five stoichiometries of assembled channels in Fig. 1 for groups of 33 channels were simulated as follows. The first step was to generate a group of 33 channels by assuming random assembly of mutant and WT subunits drawn from equal numbers of mutant and WT subunits (with replacement) for each channel in the group. To assemble each channel, four random numbers between 0 and 1 were drawn. Each random number < 0.5 indicated a WT subunit and each random number ≥ 0.5 indicated a mutant subunit. The number of channels of each of the five stoichiometries for each group of 33 channels were then binned into separate histograms for each stoichiometry. This process was then repeated for 10^6^ groups of 33 channels. Each histogram of the numbers of channels of each type for the 10^6^ groups was then normalized to a discrete probability distribution with a summed probability of 1.0, to indicate the probability of observing the indicated number of channels of that stoichiometry in a group of 33 channels. These predicted discrete probability distributions are presented in Supplementary Fig. 4 for comparison to the experimentally observed number indicated by red histogram bars in the same figure.

### Software

The computer program for simulating the discrete probability distributions for the different types of channels assuming random assembly of mutant and WT subunits, Binomdouble4-33.bas, is written in QB64 and is available from the corresponding author upon request. The increments of single channel conductance and *V*_h_ added by each G375R mutant and WT subunit for Figs. 5C and 6D were optimized by minimizing the least squared differences between the experimental values and those calculated with the equations in the legends of Figs. 5 and 6 by running Solver in Excel.

### Statistics

Error bars in the figures and error estimates in the text are SEM. Significance was determined with the two-tailed *t*-test unless otherwise indicated.

## Data Availability

The data for this study are contained in the figures and text of the manuscript and the Supplementary information.

## Acknowledgements

This work was supported in part by NIH grant R01-GM114694 to L.S. and K.L.M. We thank Julia Bulova for technical help during the initial phase of this project.

## Author contributions

Y.G., P.L., L.S., and K.L.M. designed research; Y.G., P.L., A.B., B.W., L.S., and K.L.M performed research; L.S. and K.L.M. wrote the manuscript with input from the other authors.

## Additional information

**Supplementary information** is included.

**Correspondence** and requests for materials can be addressed to Lawrence Salkoff or Karl L. Magleby.

## Supplementary information

**Supplementary Fig. 1.**
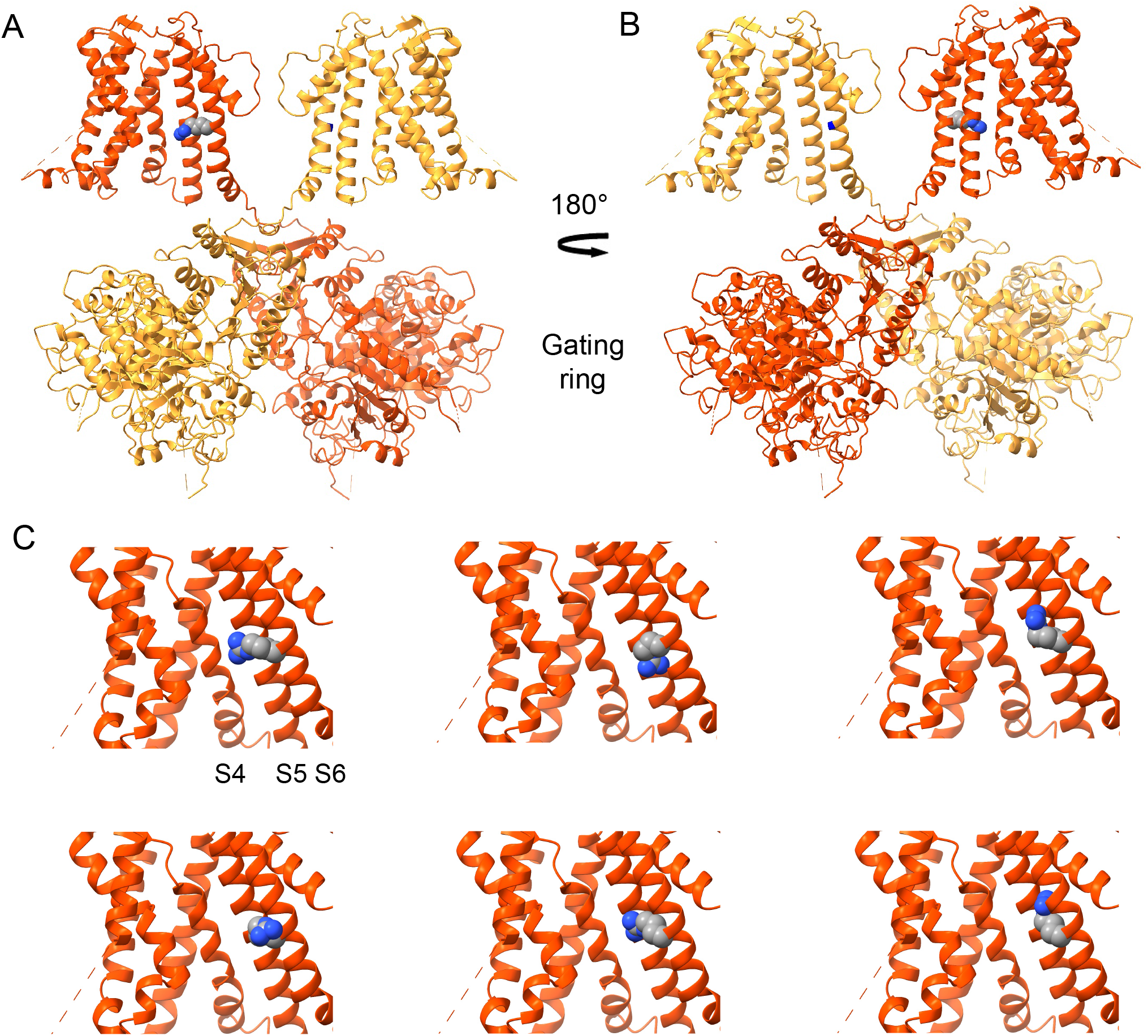
Cryo-EM ribbon structures of the Ca^2+^-free human BK channel (PDB ID:6V3G)^1^ showing the location of the mutant arginine side chain G375R. (***A***) BK channel structure with the front and back (pore-forming alpha) subunits removed. The arginine side chain of the G375R mutation is shown in space filling format on the back side of S6 away from the conduction pathway on the red colored subunit. Carbon atoms are gray, nitrogen blue and hydrogen not indicated. (***B***) Structure after rotating 180° so that the location (blue mark) of the WT glycine side chain of G375 is visible on the back side of S6 on the yellow subunit. (***C***) Six possible orientations of the mutant arginine side chain are shown, generated by ChimeraX software www.cgl.ucsf.edu/chimerax. The mutation G375D (G310D in the referenced publication) also produces a left shift in activation^2^ as G375R does, but of half the magnitute. This raises the possibility that the left shift may be related to the bulk of the substituted side chain rather than the specific charge, which is reversed for these two mutations.

**Supplementary Fig. 2.**
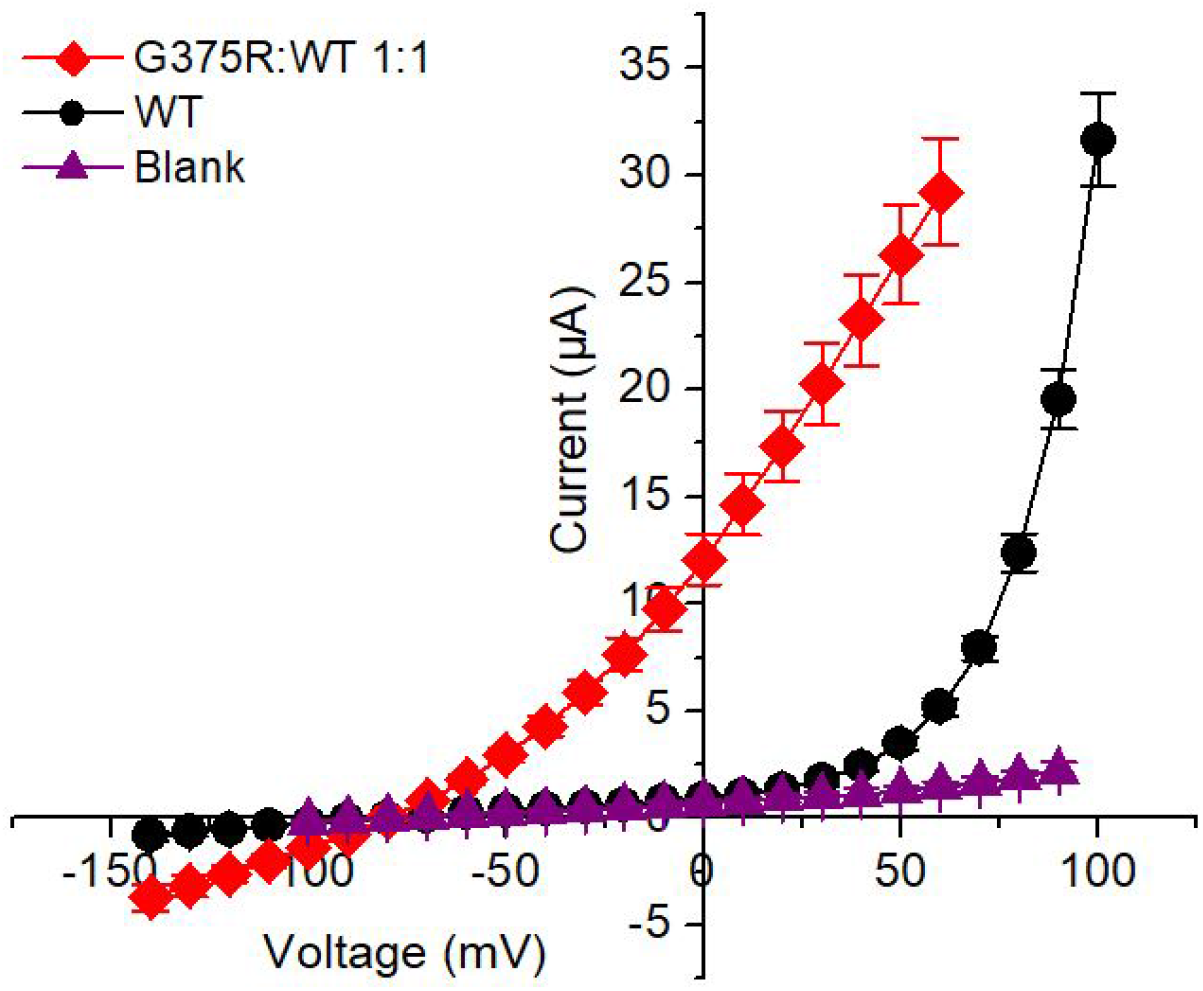
BK currents in *Xenopus* oocytes following injection of a 1:1 mixture of G375R mutant and WT cRNA activate at greatly left shifted negative voltages compared to BK currents following injection of only WT cRNA. I-V plots for the indicated injections of cRNA and blank. Two-electrode whole cell voltage clamp. These currents were used to calculate the relative conductance plotted in Fig. 2. Protocol details are in Fig. 2 legend. The amount of WT cRNA injected was 25 times greater than for the injection of the 1:1 mixture of G375R mutant and WT cRNA, so that the initial deviation of WT currents from the baseline could be readily detected.

**Supplementary Fig. 3.**
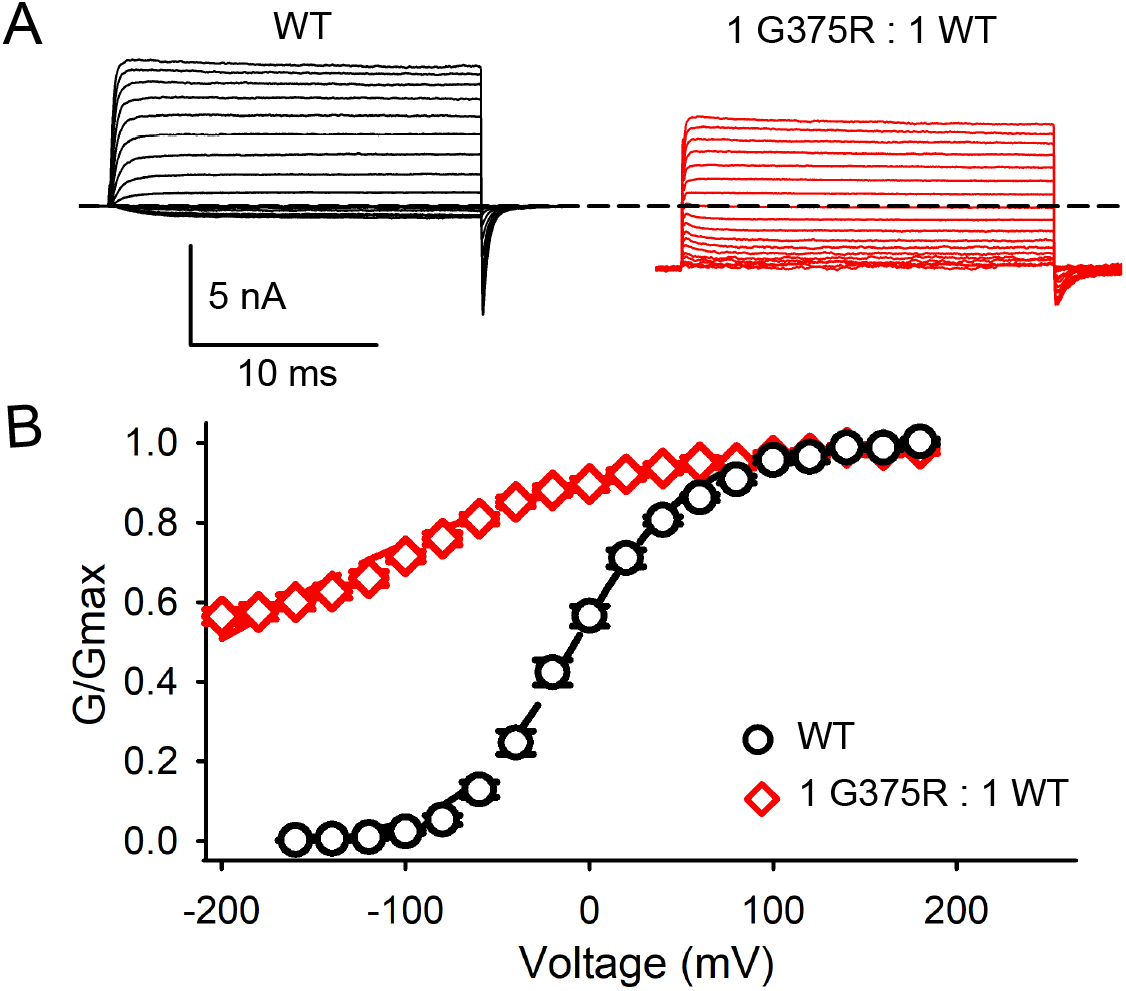
The aberrant negative shift in *V*_h_ for BK currents expressed following a 1:1 injection of G375R mutant and WT cRNA is also observed with 300 μM intracellular Ca^2+^. (***A***) Currents recorded from inside-out macro patches of membrane with 300 μM Ca^2+^ at the inner membrane surface. For WT channels voltage pulses were from −160 mV to 180 mV with 20 mV steps from a holding potential of −160 mV. Conductance was measured from tail currents after stepping to −160 mV. For channels expressed following injection of a 1:1 mixture of G375R mutant and WT cRNA, voltage pulses were from −200 mV to 180 mV with 20 mV steps from a holding potential of −200 mV. Conductance was measured from tail currents after stepping to −200 mV. The dashed line indicates the level of 0 current. In the presence of 300 μM intracellulear Ca^2+^, more than half of the channels expressed following the 1:1 injection of G375R mutant and WT cRNA remain open at −200 mV. The macro currents are the average response from many hundreds of BK channels in each macro patch. (***B***) G/Gmax vs. voltage plots following injection of WT cRNA alone (black circles) or a 1:1 injection of mutant and WT cRNA (red diamonds). The mean voltage for half activation *V*_h_ for WT currents was −7.2 ± 3.3 mV, shifting to −205 ± 9.6 mV for BK currents expressed from a 1:1 mixture of mutant and WT cRNA, producing a left shift of −198 mV with 300 μM Ca^2+^ at the inner membrane surface. This can be compared to a left shift of −120 mV with 0 Ca^2+^ (Fig. 3). Hence, the aberrant left shift in *V*_h_ induced by a 1:1 injection of mutant and WT cRNA occurs in the presence and absence of intracellular Ca^2+^.

**Supplementary Fig. 4.**
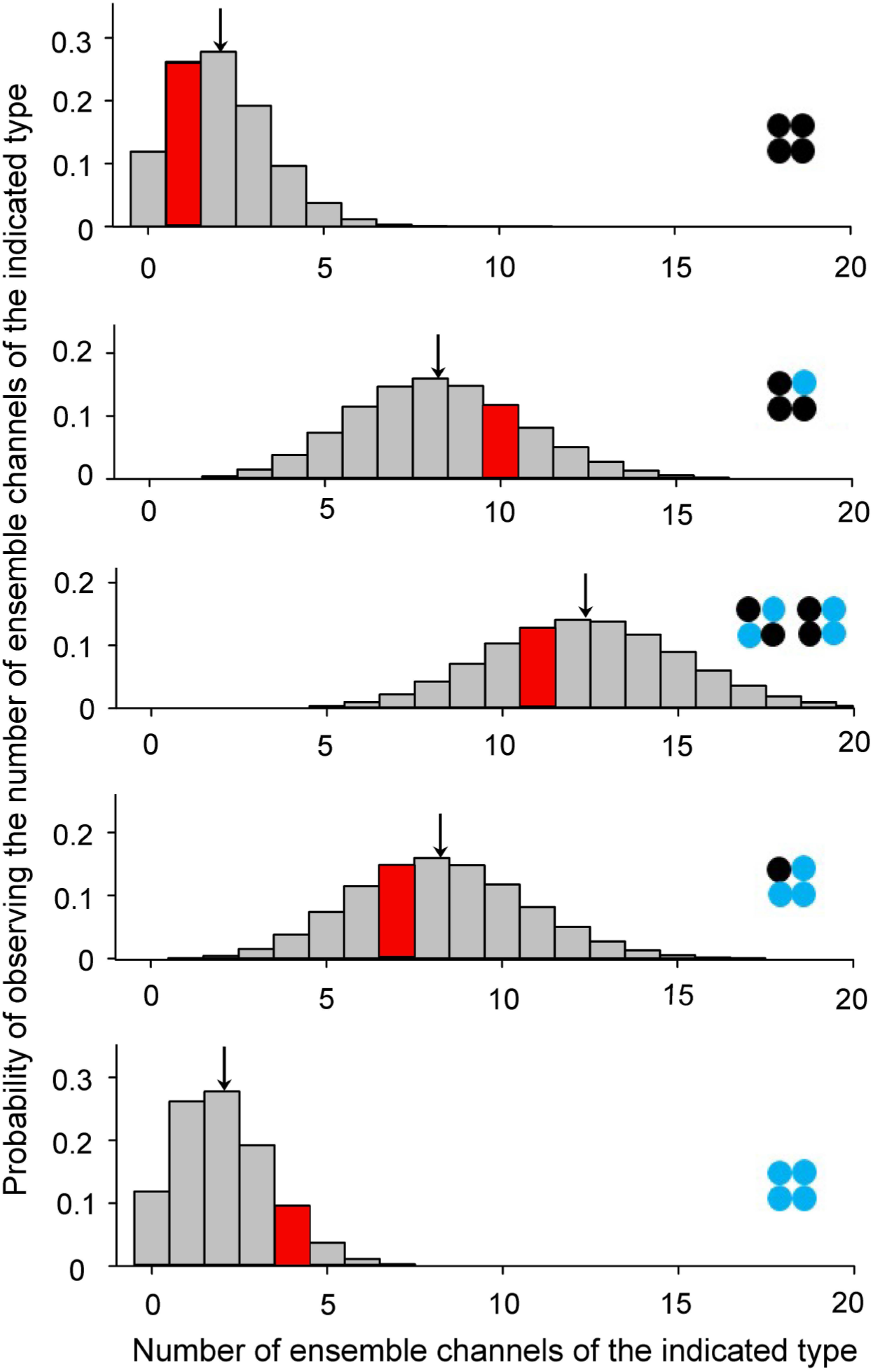
The experimentally observed percentages of the five types of assembled channels expressed following a 1:1 injection of G375R mutant and WT cRNA did not differ significantly from the theoretical percentages in Fig. 1 calculated assuming random assembly of mutant and WT subunits. To examine if the experimentally observed number of assembled channels of each type in Fig. 6 for the 33 assembled channels are consistent with random assembly of subunits, we compared the experimental number of each type to the expectations for random assembly. The expected number were determined by simulating 106 groups of 33 assembled channels and binning the number of assembled channels of each type for each group of 33 into five discrete probability distributions. The five panels in this figure present the five discrete probability distributions of the expected number of assembled channels of each type for a group of 33 channels. The channel type of each distribution is indicated by subunit composition, where black circles are WT subunits and blue circles are mutant subunits. The topmost discrete probability distribution gives the mean probabilities based on random assembly of subunits (Fig. 1) of observing the indicated number of WT channels in a group of 33 assembled channels: 11.9% of the groups of 33 assembled channels would have 0 WT assembled channels, 26.2% of the groups of 33 assembled channels would have 1 WT assembled channel (red bar), 27.8% of the groups of 33 assembled channels would have 2 WT assembled channels, and so forth. The arrow at 2.06 WT channels indicates the mean number of predicted WT channels per group of 33 assembled channels, determined from the discrete probability distribution. The mean number can also be calculated directly from the percentages in Fig. 1: 6.25% of the assembled channels are WT times 33 assembled channels equals a mean of 2.06 WT assembled channels for each group of 33. Discrete probability distributions for each of the four other types of assembled channels in Fig. 1, are also presented and are interpreted in a similar manner. For example, the discrete probability distribution for channels with 1 mutant subunit and 3 WT subunits indicates that 7.34% of the groups of 33 assembled channels would contain 5 channels of this type, 11.5% of the groups of 33 channels would contain 6 channels of this type, and so forth. Our experimental observations of the numbers of assembled channels of each type for the experimental group of 33 assembled channels (Fig. 4D and 6A, B) are indicated by the locations of the red histogram bars on the abscissa. The amplitude of the red bars gives the probability of observing the indicated number of channels of that type in the experimental group of 33 assembled channels. For example, the probability, assuming random assembly of subunits, for our observation of 1 WT channel in the experimental group of 33 assembled channels was 0.262, and the probabilities for our observations of 10, 11, 7, and 4 assembled channels for the other indicated types of assembled channels in the experimental group of 33 assembled channels were 0.118, 0.128, 0.147, and 0.0961. All of these probabilities for the observed numbers of assembled channels of each type were > 0.05 and none of the observed numbers of assembled channel types fell within the 0.05 cumulative probability of either the left or right tails of the discrete probability distributions, indicating that the experimental observations for the number of the five types of assembled channels were not significantly different^3^ from the theoretical predictions based on an assumption of random subunit assembly. A lack of significant difference does not exclude that there may be underlying differences between experimental and theoretical predictions, which could be revealed by larger sample sizes.

